# DNA Methylation Biomarkers Of Myocardial Infarction And Cardiovascular Disease

**DOI:** 10.1101/707315

**Authors:** Alba Fernández-Sanlés, Sergi Sayols-Baixeras, Isaac Subirana, Mariano Sentí, S Pérez-Fernández, Manuel Castro de Moura, Manel Esteller, Jaume Marrugat, Roberto Elosua

**Affiliations:** Cardiovascular Epidemiology and Genetics Research Group, REGICOR Study group, IMIM (Hospital del Mar Medical Research Institute), Barcelona, Catalonia, Spain; Pompeu Fabra University (UPF), Barcelona, Catalonia, Spain; Medical Research Council (MRC) Integrative Epidemiology Unit at the University of Bristol, Bristol, UK; CIBER Cardiovascular Diseases (CIBERCV), Spain; Department of Medical Sciences, Molecular Epidemiology, Uppsala University, Sweden; CIBER Epidemiology and Public Health (CIBERESP), Spain; Cancer Epigenetics and Biology Program (PEBC), Bellvitge Biomedical Research Institute (IDIBELL), Barcelona, Catalonia, Spain; CIBER Cancer (CIBERONC), Spain; Catalan Institution for Research and Advanced Studies (ICREA), Barcelona, Catalonia, Spain; Physiological Sciences Department, School of Medicine and Health Sciences, University of Barcelona (UB), Barcelona, Catalonia, Spain; Josep Carreras Leukaemia Research Institute (IJC), Badalona, Catalonia, Spain; Medicine Department, Medical School, University of Vic-Central University of Catalonia (UVic-UCC), Vic, Catalonia, Spain

**Keywords:** DNA methylation, Epigenome-wide association study, Predictive biomarkers, Myocardial infarction, Cardiovascular disease

## Abstract

**Objective:** To assess the association between DNA methylation and acute myocardial infarction, the predictive added value of the identified methylation marks, and the causality of those associations.

**Approach and Results:** We conducted a case-control, two-stage, epigenome-wide association study on acute myocardial infarction (n_discovery_=391, n_validation_=204). DNA methylation was assessed using the Infinium MethylationEPIC BeadChip (over 850,000 CpGs). DNA methylation was the exposure variable and myocardial infarction the outcome of interest. After a fixed-effects meta-analysis, 34 CpGs fulfilled Bonferroni significance. These findings were also analysed in two independent cohort studies (n∼1,800 and n∼2,500) with incident coronary (CHD) and cardiovascular disease (CVD). The Infinium HumanMethylation450 BeadChip was used in these two studies (over 480,000 CpGs) and only 12 of the 34 CpGs were available in those samples. Finally, we validated four of them in association with incident CHD: *AHRR*-mapping cg05575921, *PTCD2-*mapping cg25769469, intergenic cg21566642 and *MPO*-mapping cg04988978. The four CpGs were also associated with classical cardiovascular risk factors. A methylation risk score based on those CpGs did not improve the predictive capacity of the Framingham risk function. To assess the causal effects of those CpGs we performed Mendelian randomization analysis but only one metQTL could be identified and the results were not conclusive.

**Conclusions:** We have identified 34 CpGs related to acute myocardial infarction. These loci highlight the relevance of smoking, lipid metabolism, and inflammation in the biological mechanisms related to myocardial infarction. Four were additionally associated with incident CHD and CVD but did not provide additional predictive information.

## INTRODUCTION

Cardiovascular disease (CVD) and more specifically coronary heart disease (CHD) remains the number one cause of death and disease burden worldwide.^1,2^ At the individual level, prevention is based on the estimation of cardiovascular risk.^3^ However, the sensitivity of cardiovascular risk estimation is low and a significant proportion of CHD events occurs in individuals classified as having moderate or low risk.^4^ Additionally, the use of currently available drugs to control classical cardiovascular risk factors (CVRFs) does not prevent all CHD events, underlining the need to identify new strategies for reducing this residual cardiovascular risk.^5^ Thus, information encoded in biological mechanisms should be unravelled to find new predictive biomarkers and potential therapeutic targets. Among these biomarkers, DNA methylation marks arise as emerging candidates.

DNA methylation is an epigenetic mechanism consisting on chemical modifications of cytosines, mostly followed by guanines (CpGs).^6^ Epigenome-wide association studies (EWASs) make it possible to find DNA methylation biomarkers of different traits and outcomes. In fact, DNA methylation pattern is associated with multiple chronic diseases,^7^ including CVD and CHD.^8–13^ However, the clinical value of the identified biomarkers is still limited, and the epigenetic landscape underlying CVD is not completely understood.

The most common technology to assess DNA methylation is based on commercial arrays, which do not cover the whole methylome. Moreover, most current knowledge on the relation between DNA methylation and cardiovascular risk comes from studies based on the Infinium HumanMethylation450 BeadChip (Illumina, CA, USA; from now on, 450k)^14^ – which has been replaced by the Infinium MethylationEPIC BeadChip (Illumina, CA, USA; from now on, EPIC). Compared to the 450k, EPIC interrogates 413,745 more methylation sites (but excludes 42,859) increasing the genomic coverage. Moreover, EPIC is enriched with functional sites analyses such as enhancers, DNase hypersensitive sites, and miRNA promoter regions.^15^ Thus, the new chip has the potential to identify novel DNA methylation-based biomarkers of cardiovascular events.

We hypothesized that DNA methylation is associated with MI risk, and that some of these epigenetic marks could be predictive of future risk, and have causal effects on cardiovascular outcomes. Thus, this study had three aims: 1) to unravel genomic methylation loci associated with myocardial infarction (MI), 2) to assess their predictive capacity of cardiovascular risk, and 3) to decipher the causality of those associations.

## MATERIALS AND METHODS

### Study design and populations

We designed an epigenome-wide association study (EWAS) using three populations: the Girona Heart Registry (REGICOR, REgistre GIroní del COR), the Women’s Health Initiative (WHI), and the Framingham Offspring Study (FOS). We first performed a two-stage EWAS on acute MI using two independent age- and sex-matched case-control studies designed in REGICOR. Then, we validated the significant results in the other two populations with incident cases of CHD and CVD.

#### Case-control studies of acute MI in REGICOR

The sample used in the discovery stage (REGICOR-1) involved 416 individuals (208 MI cases and 208 controls). The sample in the validation stage (REGICOR-2) comprised 208 individuals (104 cases and 104 controls). Cases were selected from patients who were consecutively attended for a first acute MI in the reference hospital of the monitored area, in the province of Girona, in the northeast of Spain. Women were overrepresented to achieve their inclusion as 50% of our sample. Controls were participants in a population-based survey performed in the same monitored area. They were randomly selected from those attending the 2009-2013 follow-up visit (n=4,980), and matched by age and sex with the MI cases.^16^ All participants were of European descent and provided informed written consent. The study was approved by the local ethics committee (2015/6199/I; 2018/7855/I) and meets the principles expressed in the Declaration of Helsinki and the relevant Spanish legislation.

#### Samples with incident cases of CHD and CVD

The WHI sample is a case-control study nested in a cohort. The FOS sample is a prospective cohort study. Both samples were available in the database of Genotypes and Phenotypes (http://dbgap.ncbi.nlm.nih.gov; project number #9047). The graphical abstract shows the design and flow-chart of this study.

### Assessment of cardiovascular outcomes

The outcomes assessed were acute MI in REGICOR, and incident CHD and CVD in the WHI and FOS samples. Additional details are provided in the Supplementary material.

### Assessment of DNA methylation

DNA methylation was assessed genome-wide from peripheral blood with commercial arrays from Illumina (CA, USA). The Infinium MethylationEPIC BeadChip, covering over 850,000 CpGs, was used in the REGICOR samples. The Infinium HumanMethylation450 BeadChip, covering over 480,000 CpGs, was used in the WHI and FOS samples. A detailed quality control pipeline for the methylation data is available in the Supplementary material. Methylation status at each CpG was reported by β-values.^17^

### Covariates

In the REGICOR case-control studies the following covariates were considered: self-reported smoking, diabetes, hypercholesterolemia and hypertension (Supplementary material). In the WHI and FOS studies self-reported smoking and glycaemia, total and HDL cholesterol, and blood pressure measurement were considered. Moreover, we inferred the peripheral blood cell counts with the *FlowSorted.Blood.450k* R package.^18^ We also estimated two surrogate variables for unknown sources of potential technical or biological confounding using the *sva* R package.^19^

### Statistical analysis

All statistical analyses were performed using R version 3.4.0. The codes of the Singularity images are available in the repositories at https://github.com/regicor/methylation_ami/. A detailed description of the statistical methods is provided in the Supplementary material.

#### Association between DNA methylation and cardiovascular outcomes

Logistic regression was used in the analyses in the REGICOR and WHI samples, while Cox regression was used in the FOS sample. We considered the cardiovascular event (acute MI, CHD or CVD) as the outcome and DNA methylation as the exposure.

We defined three models. Model 1 was adjusted for estimated cell counts and two surrogate variables (plus age and ethnicity in the WHI sample, plus age and sex in the FOS samples). Model 2 was additionally adjusted for smoking. Model 3 was further adjusted for diabetes, hypercholesterolemia and hypertension.

In order to reduce epigenomic inflation, we corrected the coefficients, the standard errors and the *p* values using the *bacon* R package if necessary.^20^ We used coefficients and standard errors from the regression models as the input data and we set a random seed at 123. We selected those associations from the discovery stage (REGICOR-1) with a corrected *p-value*<10^−5^ for assessment in the validation stage (REGICOR-2). Moreover, we performed a fixed-effect meta-analysis of the corrected effect sizes observed in both stages, weighted by the inverse of the variance. We defined as statistically significant any association fulfilling the Bonferroni correction for multiple comparisons (0.05 divided by the number of probes analyzed). Thereafter, we studied the association of the identified CpGs with incident CHD and with CVD events in the WHI and the FOS samples, separately. The results from both samples were meta-analysed (for CHD and CVD, separately), and significance was defined accordingly to the number of probes analyzed.

#### Association between the identified CpGs and CVRFs

We analysed whether the methylation levels of the identified CpGs were associated with individual CVRFs in the four samples using multiple linear regression, and then meta-analysed the results. Significance was also defined accordingly to Bonferroni criteria.

#### Methylation risk score (MRS) and predictive capacity

We developed a weighted MRS based on the CpGs identified. We evaluated the association between this score and CVD and CHD incidence in the FOS sample, using Cox regression. All analyses were adjusted for age, sex, diabetes, smoking, systolic blood pressure, hypertensive treatment, and levels of total cholesterol and cholesterol in high-density lipoproteins (HDL-C).^21^ We also assessed the potential added predictive value of including the MRS in the Framingham risk function. We evaluated the increase in the discrimination and the reclassification.

### *In silico* functional analysis

We performed a pathway and network analysis of the genes found as differentially methylated in association with MI, using the Ingenuity Pathway Analysis (IPA) software (http://www.ingenuity.com/,QIAGEN, Redwood City, CA, USA). We uploaded the Gene symbol identifiers of those genes. The reference database underlying IPA was the Ingenuity Knowledge Base (Genes only). Only human annotations were considered. Pathway analyses were performed with IPA’s Core Analysis module. “Canonical pathways” and “Diseases and functions” terms with a *p*<0.05 after Benjamini-Hochberg multiple testing correction were defined as a statistically significant overrepresentation of input genes in a given process.

### Causality of associations between DNA methylation and cardiovascular outcomes

We performed Mendelian Randomization studies using the MR-Base platform.^22^ We selected methylation-level quantitative trait loci (meQTL) from a public database^23^ as instrumental variables, and interrogated their association with CHD using summary statistic data from a meta-analysis of GWAS assessing CHD.^24^ A more detailed description of the analysis is included in the Supplementary material.

## RESULTS

### Quality control of DNA methylation data, cardiovascular outcomes and covariates

We finally included 391 individuals (196 cases and 195 controls) in the REGICOR-1 sample, 204 individuals (101 cases and 103 controls) in the REGICOR-2 sample, 1,863 women in the WHI sample, and 2,540 participants in the FOS sample. The main sociodemographic and clinical characteristics of the three populations are shown in Tables 1 and 2. Regarding the number of CpGs, we analyzed 811,610 CpGs in the REGICOR-1 sample, 820,183 CpGs in the REGICOR-2 sample, 478,369 CpGs in the WHI sample, and 483,656 CpGs in the FOS sample.

**Table 1.** Descriptive characteristics of the populations used in the two-stage EWAS on acute myocardial infarction (AMI): REGICOR-1 and REGICOR-2.

| Variables | Discovery:<br>REGICOR-1 |  | Validation:<br>REGICOR-2 |  |
| --- | --- | --- | --- | --- |
|  | N=391 | NA‡ | N=204 | NA‡ |
| **AMI cases/controls, n (%)‡** |  |  |  |  |
| *AMI cases* | 196 (50.1) | 0 | 101 (49.5) | 0 |
| *Controls* | 195 (49.9) | 0 | 103 (50.5) | 0 |
| **Age** | 63.2 (6.94) | 0 | 61.7 (6.90) | 0 |
| *AMI cases* | 63.0 (6.96) | 0 | 61.6 (6.83) | 0 |
| *Controls* | 63.3 (6.94) | 0 | 61.7 (7.01) | 0 |
| **Sex, female, n (%)** | 201 (51.4) | 0 | 100 (49.0) | 0 |
| *AMI cases* | 100 (51.0) | 0 | 49 (48.5) | 0 |
| *Controls* | 101 (51.8) | 0 | 51 (49.5) | 0 |
| **Smokers, n (%)** | 93 (24.4) | 10 | 67 (33.3) | 3 |
| *AMI cases* | 76 (40.6) | 9 | 45 (45.0) | 1 |
| *Controls* | 17 (8.76) | 1 | 22 (21.8) | 2 |
| **BMI, kg/m2\*‡** | 28.5 (4.75) | 52 | 27.5 (4.87) | 23 |
| *AMI cases* | 28.1 (4.34) | 52 | 28.2 (5.87) | 23 |
| *Controls* | 28.8 (5.02) | 0 | 26.9 (3.90) | 0 |
| **Hypercholesterolaemia, n (%)‡** | 179 (53.0) | 53 | 93 (51.4) | 23 |
| *AMI cases* | 93 (64.1) | 51 | 51 (65.4) | 23 |
| *Controls* | 86 (44.6) | 2 | 42 (40.8) | 0 |
| **HTN, n (%)‡** | 212 (57.1) | 20 | 105 (54.1) | 10 |
| *AMI cases* | 120 (68.2) | 20 | 62 (68.1) | 10 |
| *Controls* | 92 (47.2) | 0 | 43 (41.7) | 0 |
| **Diabetes, n (%)** | 87 (24.7) | 39 | 44 (23.8) | 19 |
| *AMI cases* | 56 (35.7) | 39 | 27 (32.9) | 19 |
| *Controls* | 31 (15.9) | 0 | 17 (16.5) | 0 |

\*Mean (Standard deviation).
‡Median (Interquartile Range).
‡ NA, missing information; AMI, Acute myocardial infarction; BMI, Body mass index; Hypercholesterolaemia, defined as self-reported high cholesterol levels or treatment; HTN, Hypertension, defined as self-reported high blood pressure or treatment; Diabetes, defined as self-reported diabetes or treatment.

**Table 2.** Descriptive characteristics of the populations used in the follow-up association studies on incident cardiovascular (CVD) and coronary heart disease (CHD) events: Women’s Health Initiative (WHI) and Framingham Offspring Study (FOS).

| Variables | WHI | NA | FOS CHD | NA | FOS CVD | NA |
| --- | --- | --- | --- | --- | --- | --- |
|  | N=1863 |  | N=2266 |  | N=2123 |  |
| Age | 64.2 (7.03) | 0 | 65.7 (8.82) | 0 | 65.3 (8.66) | 0 |
| Sex, female, n (%) | 1863 (100) | 0 | 1284 (56.7) | 0 | 1206 (56.8) | 0 |
| CHD, n (%)‡ | 914 (49.1) | 0 | 106 (4.68) | 0 | — | — |
| CVD, n (%)‡ | 984 (52.8) | 0 | — | — | 222 (10.5) | 0 |
| Smokers, n (%) | 174 (9.46) | 24 | 217 (9.62) | 11 | 202 (9.56) | 11 |
| BMI, kg/m2*‡ | 29.8 (5.97) | 11 | 28.1 (5.39) | 8 | 28.0 (5.33) | 6 |
| Total cholesterol, mg/dl* | 233 (42.3) | 1 | 189 (36.2) | 1 | 190 (35.8) | 1 |
| LDL-C, mg/dl*‡ | 152 (38.7) | 31 | 107 (30.8) | 2 | 108 (30.5) | 2 |
| HDL-C, mg/dl*‡ | 52.2 (13.2) | 1 | 58.3 (18.3) | 2 | 58.7 (18.3) | 2 |
| TG, mg/dl† | 126 [93.0;176] | 1 | 101 [73.0;140] | 1 | 100 [73.0;139] | 1 |
| Cholesterol treatment, n (%) | 166 (11.4) | 403 | 871 (38.5) | 3 | 787 (37.1) | 3 |
| Hypercholesterolaemia, n (%)‡ | 913 (55.8%) | 227 | 1033 (45.7) | 4 | 943 (44.5) | 4 |
| SBP, mmHg*‡ | 131 (17.5) | 0 | 126 (16.9) | 1 | 125 (16.7) | 1 |
| DBP, mmHg*‡ | 76.4 (9.27) | 0 | 72.1 (9.99) | 3 | 72.3 (9.92) | 3 |
| HTN, n (%)‡ | 952 (61.2) | 308 | 1217 (53.9) | 8 | 1110 (52.5) | 7 |
| HTN treatment, n (%)‡ | 672 (45.9) | 400 | 1025 (45.4) | 7 | 927 (43.8) | 6 |
| Glucose, mg/dl† | 107 (38.3) | 1 | 106 (21.8) | 2 | 105 (21.5) | 2 |
| Diabetes, n (%) | 319 (17.1) | 0 | 307 (14.3) | 126 | 266 (13.3) | 119 |
| Diabetes treatment, n (%) | 178 (9.57) | 3 | 87 (4.04) | 110 | 72 (3.56) | 102 |

\*Mean (Standard deviation). †Median (Interquartile range). ‡ CHD, Coronary heart disease; CVD, Cardiovascular disease; BMI, Body mass index; LDL-C, Cholesterol in low-density lipoprotein; HDL-C, Cholesterol in high-density lipoprotein; TG, Triglycerides; Hypercholesterolaemia, defined as previous treatment or total cholesterol $\geq 240$ mmHg or LDL-C $\geq 160$ mmHg; SBP, Systolic blood pressure; DBP, Diastolic blood pressure; HTN, Hypertension, defined as previous treatment or SBP $\geq 140$ mmHg or DBP $\geq 90$ mmHg; Diabetes, defined as previous treatment or glycaemia $\geq 126$ mg/dl.

### Association between DNA methylation and cardiovascular outcomes

#### A. Two-stage EWAS on acute myocardial infarction

##### A.1. Discovery stage

The associations from the discovery stage (REGICOR-1) that were taken to the subsequent validation (*p-value*<10^−5^), and their Manhattan and Q-Q plots are shown in the Supplementary Table 1, and Supplementary Figures 1-2. In total, we identified 68 suggestively significant CpGs (Supplementary Figure 3). Model 1 provided 56 CpGs, of which three were also found in both model 2 and 3, and 13 in model 2. One additional CpG was found in both model 2 and 3, two in model 2 and nine in model 3.

##### A.2. Validation and meta-analysis

The association studies performed in the validation stage included the 68 suggestively significant CpGs. We meta-analyzed the results of those 68 associations from both stages. We identified 34 differentially methylated CpGs related to MI, with similar effect sizes in all three models and both stages for most of the CpGs. Specifically, all of them were statistically significant (*p-value*<6.17·10^−8^) in model 1, 17 in model 2, and three in model 3 (Table 3, and Supplementary Table 2). The 34 CpGs were located in 25 different loci (26 genes, with one CpG mapping to two genes) and nine intergenic regions. Fourteen of these CpGs should be interpreted with caution, as the *p-value* in the validation population was not significant considering the 68 CpGs analyzed (*p-value*>0.05/68=7.35·10^−4^) (Table 3, and Supplementary Table 2).

**Table 3.**
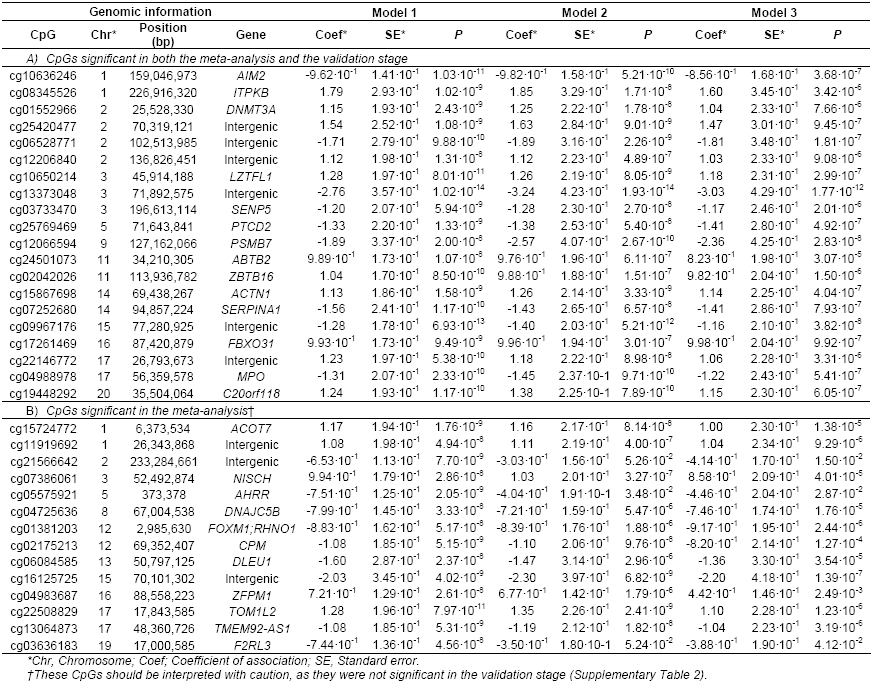
Significant CpGs differentially methylated in association with prevalent myocardial infarction in the fixed-effects meta-analyses of the REGICOR case-control samples. Model 1 was adjusted for estimated cell counts and two surrogate variables. Model 2 was further adjusted for smoking status. Model 3 was additionally adjusted for diabetes, hypercholesterolemia, and hypertension. The Bonferroni-corrected *p* threshold was 1.67·10^−8^.

##### A.3. *In silico* functional analysis

We performed an IPA analysis of the 26 identified genes. Among the 10 most significant molecular and cellular functions, this set of genes was enriched on lipid metabolism. The enriched physiological system development and functions included the connective tissue, hematological system, hematopoiesis, and the endocrine system. Inflammatory response and disease, metabolic disease, and cardiovascular disease were also among the 20 most significant diseases/disorders (Supplementary Figure 4).

#### B. Follow-up association studies on incident CHD and CVD events

Out of the 34 identified CpGs associated with MI, only 12 were available in the samples with incident cases (whose DNA methylation was profiled with the array 450k). In total, we validated four CpGs after the meta-analysis of the separate association studies in the WHI and the FOS samples (*p-value*<0.05/12=4.17·10^−3^): *AHRR*-mapping cg05575921, *PTCD2-* mapping cg25769469, intergenic cg21566642 and *MPO*-mapping cg04988978. The four CpGs were associated with CHD but cg25769469 was not related to CVD (Table 4, Supplementary Table 3).

**Table 4.**
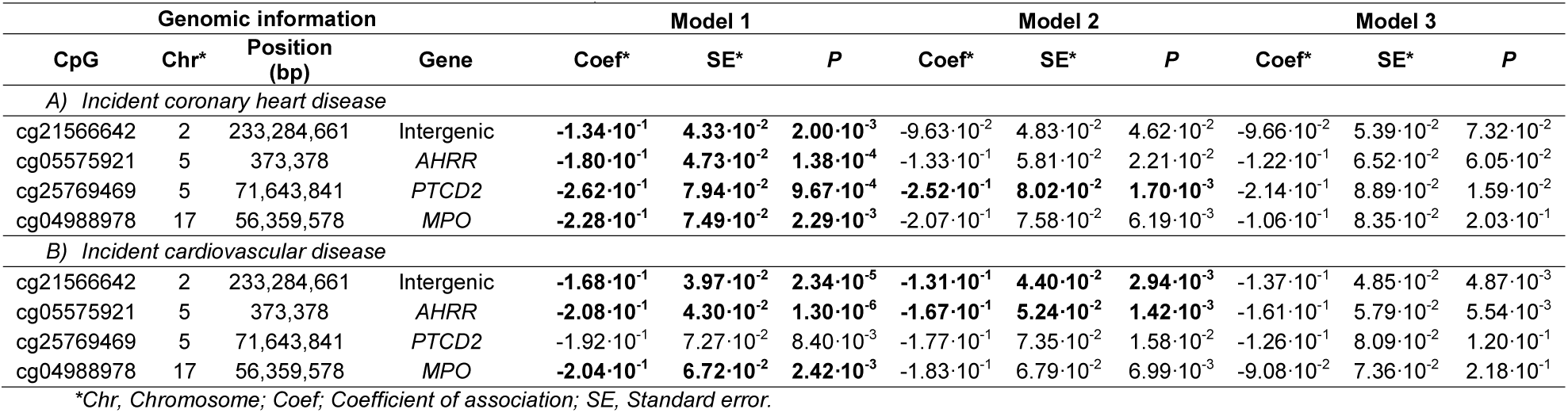
Significant CpGs differentially methylated in association with incident cardiovascular/coronary disease in the fixed-effects meta-analyses of the samples from the Women’s Health Initiative (WHI) and the Framingham Offspring Study (FOS). Model 1 was adjusted for age, estimated cell counts and two surrogate variables (plus ancestry in WHI, plus sex in FOS). Model 2 was further adjusted for smoking status. Model 3 was additionally adjusted for diabetes, hypercholesterolemia, and hypertension. The Bonferroni-corrected *p* threshold was 4.17·10^−3^ (those significant are highlighted in bold).

| Genomic information |  |  |  | Model 1 |  |  | Model 2 |  |  | Model 3 |  |  |
| --- | --- | --- | --- | --- | --- | --- | --- | --- | --- | --- | --- | --- |
| CpG | Chr* | Position (bp) | Gene | Coef* | SE* | P | Coef* | SE* | P | Coef* | SE* | P |
| A) Incident coronary heart disease |  |  |  |  |  |  |  |  |  |  |  |  |
| cg21566642 | 2 | 233,284,661 | Intergenic | **-1.34·10<sup>-1</sup>** | **4.33·10<sup>-2</sup>** | **2.00·10<sup>-3</sup>** | -9.63·10<sup>-2</sup> | 4.83·10<sup>-2</sup> | 4.62·10<sup>-2</sup> | -9.66·10<sup>-2</sup> | 5.39·10<sup>-2</sup> | 7.32·10<sup>-2</sup> |
| cg05575921 | 5 | 373,378 | AHRR | **-1.80·10<sup>-1</sup>** | **4.73·10<sup>-2</sup>** | **1.38·10<sup>-4</sup>** | -1.33·10<sup>-1</sup> | 5.81·10<sup>-2</sup> | 2.21·10<sup>-2</sup> | -1.22·10<sup>-1</sup> | 6.52·10<sup>-2</sup> | 6.05·10<sup>-2</sup> |
| cg25769469 | 5 | 71,643,841 | PTCD2 | **-2.62·10<sup>-1</sup>** | **7.94·10<sup>-2</sup>** | **9.67·10<sup>-4</sup>** | **-2.52·10<sup>-1</sup>** | **8.02·10<sup>-2</sup>** | **1.70·10<sup>-3</sup>** | -2.14·10<sup>-1</sup> | 8.89·10<sup>-2</sup> | 1.59·10<sup>-2</sup> |
| cg04988978 | 17 | 56,359,578 | MPO | **-2.28·10<sup>-1</sup>** | **7.49·10<sup>-2</sup>** | **2.29·10<sup>-3</sup>** | -2.07·10<sup>-1</sup> | 7.58·10<sup>-2</sup> | 6.19·10<sup>-3</sup> | -1.06·10<sup>-1</sup> | 8.35·10<sup>-2</sup> | 2.03·10<sup>-1</sup> |
| B) Incident cardiovascular disease |  |  |  |  |  |  |  |  |  |  |  |  |
| cg21566642 | 2 | 233,284,661 | Intergenic | **-1.68·10<sup>-1</sup>** | **3.97·10<sup>-2</sup>** | **2.34·10<sup>-5</sup>** | **-1.31·10<sup>-1</sup>** | **4.40·10<sup>-2</sup>** | **2.94·10<sup>-3</sup>** | -1.37·10<sup>-1</sup> | 4.85·10<sup>-2</sup> | 4.87·10<sup>-3</sup> |
| cg05575921 | 5 | 373,378 | AHRR | **-2.08·10<sup>-1</sup>** | **4.30·10<sup>-2</sup>** | **1.30·10<sup>-6</sup>** | **-1.67·10<sup>-1</sup>** | **5.24·10<sup>-2</sup>** | **1.42·10<sup>-3</sup>** | -1.61·10<sup>-1</sup> | 5.79·10<sup>-2</sup> | 5.54·10<sup>-3</sup> |
| cg25769469 | 5 | 71,643,841 | PTCD2 | -1.92·10<sup>-1</sup> | 7.27·10<sup>-2</sup> | 8.40·10<sup>-3</sup> | -1.77·10<sup>-1</sup> | 7.35·10<sup>-2</sup> | 1.58·10<sup>-2</sup> | -1.26·10<sup>-1</sup> | 8.09·10<sup>-2</sup> | 1.20·10<sup>-1</sup> |
| cg04988978 | 17 | 56,359,578 | MPO | **-2.04·10<sup>-1</sup>** | **6.72·10<sup>-2</sup>** | **2.42·10<sup>-3</sup>** | -1.83·10<sup>-1</sup> | 6.79·10<sup>-2</sup> | 6.99·10<sup>-3</sup> | -9.08·10<sup>-2</sup> | 7.36·10<sup>-2</sup> | 2.18·10<sup>-1</sup> |

\*Chr, Chromosome; Coef; Coefficient of association; SE, Standard error.

### Association between the identified CpGs and CVRFs

Table 5 shows the associations observed between the identified CpGs and classical CVRFs. The four identified CpGs were related to some CVRF [*p-value*<0.05/(4 CpGs*8 CVRF)=1.56·10^−3^].

**Table 5.** Significant associations between the identified CpGs and classical cardiovascular risk factors (CVRFs) in the fixed-effects meta-analyses of the four samples. The significant associations are highlighted in bold (*p* < 1.56·10^−3^ †).

| CpG | Smoking | BMI* | HDL-C* | LDL-C* | TG* | Glucose | SBP | DBP* |  |
| --- | --- | --- | --- | --- | --- | --- | --- | --- | --- |
| cg21566642 | -1.61 | $9.55 \cdot 10^{-3}$ | $7.69 \cdot 10^{-4}$ | $-1.10 \cdot 10^{-3}$ | $-4.17 \cdot 10^{-4}$ | $2.32 \cdot 10^{-4}$ | $2.30 \cdot 10^{-3}$ | $7.97 \cdot 10^{-3}$ | Coefficient |
| | $4.22 \cdot 10^{-2}$ | $2.65 \cdot 10^{-3}$ | $1.21 \cdot 10^{-3}$ | $4.09 \cdot 10^{-4}$ | $1.95 \cdot 10^{-4}$ | $4.20 \cdot 10^{-4}$ | $9.02 \cdot 10^{-4}$ | $1.67 \cdot 10^{-3}$ | SE* |
| | **$<2.2 \cdot 10^{-16}$** | **$3.21 \cdot 10^{-4}$** | $5.24 \cdot 10^{-1}$ | $7.11 \cdot 10^{-3}$ | $3.22 \cdot 10^{-2}$ | $5.81 \cdot 10^{-1}$ | $1.08 \cdot 10^{-2}$ | **$1.78 \cdot 10^{-6}$** | P |
| cg05575921 | -1.61 | $1.17 \cdot 10^{-2}$ | $4.36 \cdot 10^{-3}$ | $-1.35 \cdot 10^{-3}$ | $-7.09 \cdot 10^{-4}$ | $1.86 \cdot 10^{-4}$ | $2.15 \cdot 10^{-3}$ | $7.54 \cdot 10^{-3}$ | Coefficient |
| | $4.22 \cdot 10^{-2}$ | $2.46 \cdot 10^{-3}$ | $1.11 \cdot 10^{-3}$ | $3.77 \cdot 10^{-4}$ | $1.79 \cdot 10^{-4}$ | $3.87 \cdot 10^{-4}$ | $8.23 \cdot 10^{-4}$ | $1.52 \cdot 10^{-3}$ | SE* |
| | **$<2.2 \cdot 10^{-16}$** | **$2.22 \cdot 10^{-6}$** | **$8.22 \cdot 10^{-5}$** | **$3.39 \cdot 10^{-4}$** | **$7.63 \cdot 10^{-5}$** | $6.30 \cdot 10^{-1}$ | $9.06 \cdot 10^{-3}$ | **$7.69 \cdot 10^{-7}$** | P |
| cg25769469 | $-9.01 \cdot 10^{-2}$ | $-1.12 \cdot 10^{-2}$ | $6.76 \cdot 10^{-3}$ | $2.47 \cdot 10^{-4}$ | $-1.38 \cdot 10^{-3}$ | $-1.52 \cdot 10^{-3}$ | $-2.35 \cdot 10^{-3}$ | $-4.17 \cdot 10^{-3}$ | Coefficient |
| | $5.08 \cdot 10^{-2}$ | $2.64 \cdot 10^{-3}$ | $1.20 \cdot 10^{-3}$ | $4.11 \cdot 10^{-4}$ | $1.93 \cdot 10^{-4}$ | $4.17 \cdot 10^{-4}$ | $9.10 \cdot 10^{-4}$ | $1.69 \cdot 10^{-3}$ | SE* |
| | $7.65 \cdot 10^{-2}$ | **$2.32 \cdot 10^{-5}$** | **$1.92 \cdot 10^{-8}$** | $5.48 \cdot 10^{-1}$ | **$8.75 \cdot 10^{-13}$** | **$2.69 \cdot 10^{-4}$** | $9.97 \cdot 10^{-3}$ | $1.36 \cdot 10^{-2}$ | P |
| cg04988978 | $-7.29 \cdot 10^{-2}$ | $-1.15 \cdot 10^{-2}$ | $5.28 \cdot 10^{-3}$ | $3.59 \cdot 10^{-4}$ | $-1.19 \cdot 10^{-3}$ | $-1.88 \cdot 10^{-3}$ | $-2.88 \cdot 10^{-3}$ | $-3.54 \cdot 10^{-3}$ | Coefficient |
| | $5.15 \cdot 10^{-2}$ | $2.64 \cdot 10^{-3}$ | $1.20 \cdot 10^{-3}$ | $4.08 \cdot 10^{-4}$ | $1.92 \cdot 10^{-4}$ | $4.15 \cdot 10^{-4}$ | $9.02 \cdot 10^{-4}$ | $1.67 \cdot 10^{-3}$ | SE* |
| | $1.57 \cdot 10^{-1}$ | **$1.26 \cdot 10^{-5}$** | **$1.08 \cdot 10^{-5}$** | $3.80 \cdot 10^{-1}$ | **$5.79 \cdot 10^{-10}$** | **$5.97 \cdot 10^{-6}$** | **$1.41 \cdot 10^{-3}$** | $3.47 \cdot 10^{-2}$ | P |

\*BMI, Body mass index; HDL-C, Cholesterol in high-density lipoprotein; LDL-C, Cholesterol in low-density lipoprotein; TG, Triglycerides; SBP, Systolic blood pressure; DBP, Diastolic blood pressure; SE, Standard Error. † $0.05/(\text{number of CpGs} \cdot 8 \text{ CVRFs})$

### Association between MRS and incidence of CHD and CVD

The associations between MRS and the incidence of coronary (n=94) and cardiovascular (n=222) events in the FOS population are shown in Supplementary Table 4. The median of the follow-up periods for CVD and CHD incidence were 7.67 and 7.87 years, respectively. The MRS was not associated with higher cardiovascular risk independently of the classical CVRFs. Consistently, the addition of the MRS to the Framingham risk function did not improve its predictive capacity in the FOS cohort (Supplementary Table 4).

### Causality of the associations between DNA methylation and cardiovascular outcomes

We identified one genetic variant (rs72617176) associated with cg21566642 methylation level. Only the Wald ratio method could be performed, and the association was not significant (Supplementary Table 5). We couldn’t create genetic instrumental variables to assess the causality of the association of the other three CpGs with CHD.

## DISCUSSION

We have identified 34 methylation loci associated with acute MI in a two-stage EWAS, analyzing ∼850,000 CpGs. All but two of these MI-associated sites (one located in *AHRR* and the other one intergenic) are newly reported. Of those, 12 CpGs could be studied in association with incident cases of CHD and CVD, and we identified four of them associated with incident CHD (three of them also with incident CVD). All four were also related to traditional CVRFs, supporting their role in the development of these diseases. However, their clinical utility as predictive biomarkers or drug targets was not proven.

Recently, two EWASs on incident CHD were published providing different findings from ours. Ward-Caviness *et al* found nine CpGs associated with incident acute MI.^9^ Agha *et al* reported 52 CpGs related to incident CHD.^8^ None of them was replicated in our study. This lack of concordance could be related to methodological differences (incident vs prevalent cases; considered confounder variables), and highlights the complexity of the study of these diseases.

### CpG sites associated with acute MI events

The 34 identified CpGs showed similar effect sizes in the two REGICOR samples. Although 14 CpGs were non-significant in the validation sample, we considered them potentially relevant because the magnitude of their association with MI was comparable across the two samples and the statistical power of the validation sample was lower. Similarly, all but three CpGs (*AHRR*-mapping cg05575921, *F2RL3*-mapping cg03636183, and the intergenic cg21566642) showed consistent effect sizes in the three models. The effect size of those three was reduced by half when adjusted for smoking, which highlights the important role of this risk factor in the MI context. In fact, all three sites are widely described to be related to smoking.^16,25,26^

Differentially methylated genes were enriched in diverse molecular and physiological pathways, including lipid metabolism and metabolic and inflammatory diseases, underlining their relevance on the pathogenesis of CHD. Considering the Bonferroni correction, three different CpGs were associated with MI independently of traditional CVRFs (smoking, diabetes, hypertension and hypercholesterolemia), suggesting that other biological pathways are triggered in the MI context. To our knowledge, no EWAS on any trait has previously found any of these three CpGs. For instance, cg12066594 is located within a regulatory region of *PSMB7*, which encodes a subunit of the proteasome, a complex essential for protein degradation and involved in inflammation.^27^ A polymorphism in a gene encoding another proteasome subunit (*PSMA6*) has been associated with CHD and MI.^28–31^ According to the EPIC manifest, the probes of cg12066594 and cg09967176 contain both of them three different genetic variants (rs114673478, rs569238855, rs539730482 in that of cg12066594; and rs570492126, rs534702669, rs181647444 in that of cg09967176). rs539730482, located in the single base extension site (one basepair upstream of the CpG site), and rs181647444, in the CpG site, are variants in an African population and their frequency is very low (Minor Allele Frequency <1%).^32,33^ Thus, as the individuals in this study are of European ancestry and these variants showed very low frequency in this population, the identified associations most likely are not confounded by the genetic variants.

Nonetheless, we cannot infer the biological sequence of the epigenetic marks, the related biological mechanisms, and the clinical event. One possible scenario could be that the identified DNA methylation marks occurred before the acute event, as potential biological mechanisms involved in MI pathogenesis. This may be the case of the three CpGs that were related to smoking. Conversely, as blood samples of MI cases were collected within the initial 24 h after hospitalization, the other possibility could be that methylation at the identified CpGs had changed as a consequence of the acute event or the therapeutic procedures. If the first scenario can be proven in further studies, these DNA methylation marks could be potential predictive biomarkers of MI or new therapeutic targets. If they are found to be post-MI marks, further studies could evaluate their potential as biomarkers of prognosis.

### CpG sites consistently related to prevalent and incident CVD events

Twelve of the 34 identified CpGs could be evaluated in prospective samples and four of them were also significantly related to incident cases of CHD. cg21566642 maps to an intergenic region, and cg05575921, cg04988978 and cg25769469 annotate to *AHRR, MPO* and *PTCD2*, respectively. To our knowledge, these CpGs were not associated with cardiovascular events in previous EWAS reports.

cg21566642 and cg05575921 were highly and inversely associated with smoking, which is supported by previous EWAS.^25,26^ We have also previously reported both CpGs as related to age-independent cardiovascular risk,^13^ and they have been related to all-cause mortality in an EWAS.^34^ cg05575921 was further associated directly with HDL-C and inversely with cholesterol in low-density lipoproteins (LDL-C) and triglyceride levels in our study. This CpG has been related to both CHD prevalence and incidence in a candidate gene study.^35^ cg04988978 and cg25769469 annotate to *MPO* and *PTCD2*, respectively. Both CpGs were associated directly with HDL-C and inversely with triglyceride and glucose levels. *MPO* encodes the myeloperoxidase, which promotes atherosclerotic lesions by enhancing APOB oxidation within low-density lipoproteins^36^ and was causally associated with incident cardiovascular outcomes.^37^ One CpG located within *PTCD2* was previously identified to be associated with hypertension in obstructive sleep apnea patients.^38^

### MRSs as predictive CVD biomarkers

To assess the value of the four identified CpGs as predictive biomarkers, we followed the AHA recommendations.^39^ However, neither we observed an association between the MRS and the incidence of CVD events in the FOS, nor we observed an improvement in the predictive capacity of the Framingham risk function when including this score. This highlights the challenge of novel biomarkers to improve cardiovascular risk prediction.

### Causality of the associations between methylation loci and cardiovascular outcomes

MR studies could only be performed on the association of cg21566642 with acute MI and CHD. Although a non-causal relationship was suggested, this must be interpreted with caution as only one genetic variable was used as the genetic instrument. In fact, the four CpGs associated not only with acute MI, but also incident CHD, may suggest that DNA methylation changes at those loci occur prior to the event.

### Strengths and limitations

The main strength of our study is that it is the first two-stage EWAS on MI to be based on more than 800,000 CpGs across the genome. Moreover, we aimed to validate our findings in prospective samples of CHD and CVD as a proxy of MI. Also, we aimed to prove the clinical relevance of our findings. However, some limitations should be acknowledged. First, two thirds of the CpGs identified in the initial case-control study could not be assessed in the incident studies as the methylation arrays differed in the number of CpGs (EPIC VS 450k, respectively). Second, we used self-reported information about cardiovascular risk factors in the case-control study, as an event such as MI modifies risk factor levels during the acute phase. Third, we cannot infer causality since changes in methylation could have occurred as a consequence of the acute phase and disease management of the MI event. We aimed to perform MR studies of the association between the identified CpGs and cardiovascular events, but available meQTL datasets are still limited. Last, our study is based on populations of European origin and the results cannot be extrapolated to other populations.

## CONCLUSIONS

Our study provides 34 novel DNA methylation loci related to MI. The results shed some light on the molecular landscape of MI, highlighting the importance of traditional CVRFs and inflammation in the development of CHD. Our results question the relevance of DNA methylation as a predictive biomarker.

## Abbreviations

450k: Infinium HumanMethylation450 BeadChip
MI: Myocardial Infarction
CHD: Coronary Heart Disease
CVD: Cardiovascular Disease
CVR: Cardiovascular Risk
CVRF: Cardiovascular Risk Factor
EPIC: Infinium MethylationEPIC BeadChip
EWAS: Epigenome-Wide Association Study
FOS: Framingham Offspring Study
MRS: Methylation Risk Score
REGICOR: REgistre GIroní del COR
WHI: Women’s Health Iniciative

## Acknowledgements

a. **Acknowledgements:** We thank Elaine M. Lilly, PhD, for her critical reading and revision of the English text.
b. **Sources of funding:** This project was funded by the Carlos III Health Institute– European Regional Development Fund (FIS PI18/00017, FIS PI15/00051, PI12/00232, CIBERCV, CIBERESP, CIBERONC), PERIS from Agència de Gestió d’Ajuts Universitaris i de Recerca (SLT002/16/00088) and the Government of Catalonia through the Agency for Management of University and Research Grants (2014SGR240, 2017SGR946). S. Sayols-Baixeras was funded by the Instituto de Salud Carlos III Fondos FEDER (IFI14/00007). A. Fernández-Sanlés was funded by the Spanish Ministry of Economy and Competitiveness (BES-2014–069718). The Framingham Heart Study (FHS) is conducted and supported by the National Heart, Lung, and Blood Institute (NHLBI) in collaboration with Boston University (Contract No. N01-HC-25195 and HHSN268201500001I). The Women’s Health Initiative (WHI) program is funded by the NHLBI (Contracts N01WH22110, 24152, 32100-2, 32105-6, 32108-9, 32111-13, 32115, 32118-32119, 32122, 42107-26, 42129-32, and 44221). This manuscript was not prepared in collaboration with investigators of the FHS/ WHI, has not been reviewed and/or approved by the FHS/WHI, and does not necessarily reflect the opinions or views of the FHS and WHI investigators or the NHLBI.

## HIGHLIGHTS

- 34 novel loci differentially methylated in relation to acute myocardial infarction were identified.
- Four of those methylation loci were further associated with incident cases of coronary heart disease.
- The identified loci highlight the relevance of traditional cardiovascular risk factors and biological mechanisms such as lipid metabolism and inflammation underlying myocardial infarction.

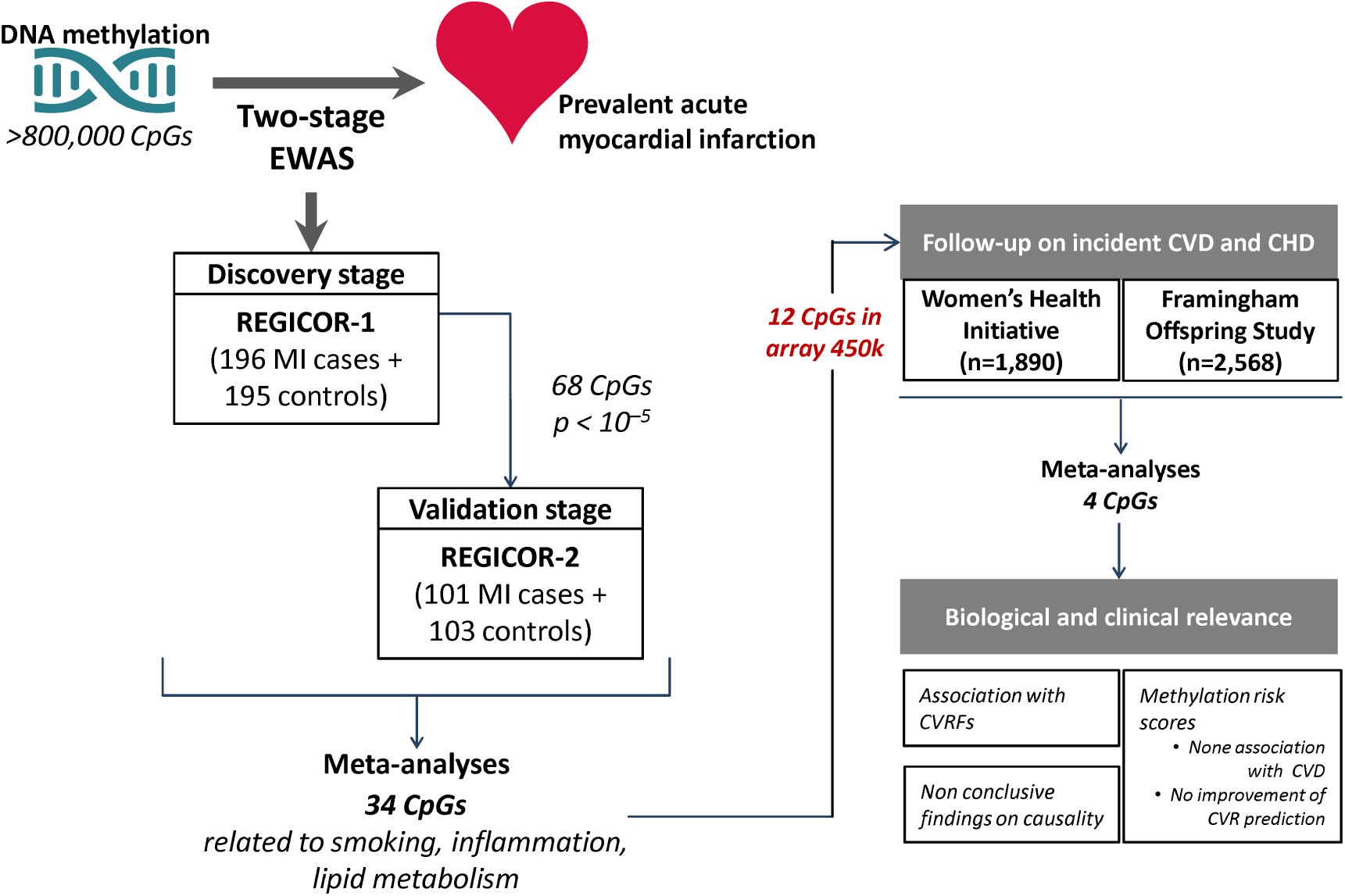

