## Supplemental file for "DNA Methylation Biomarkers Of Myocardial Infarction And Cardiovascular Disease"

**SUPPLEMENTARY MATERIAL**

**SUPPLEMENTARY METHODS**

**Assessment of cardiovascular outcomes**

*a.-* *REGICOR:* Acute myocardial infarction (AMI) was defined according to international World Health Organization criteria^1^ when two or more of the following were present: abnormal new Q waves, increase in cardiac enzymes beyond twice the upper normal value, and chest pain lasting more than 20 min.

*b.-* *WHI*: coronary heart disease (CHD) was considered when the participant had experienced any of the following: clinical AMI, definite silent AMI, total AMI, angina, coronary revascularization procedure or CHD death. Cardiovascular disease (CVD) included CHD and stroke or death due to cerebrovascular disease or other CVD. Clinical events were adjudicated in 2016 using the form 121, which considered events requiring hospitalization or occurring during a hospitalization for another reason. Diagnosis of a stroke was completed by a stroke neurologist in form 132. The cause of death was recorded using form 124.

c.- *FOS*: data regarding CHD and CVD were extracted from follow-up for cardiovascular events through 2014 (until exam 12). We used exam 8 as the baseline visit and to identify prevalent cases.

**Assessment of DNA methylation status**

In the REGICOR study, DNA was extracted from whole peripheral blood drawn within the first 24 hours post-AMI for cases and at follow-up for controls. For the other samples, DNA was extracted from buffy coat obtained from whole peripheral blood samples collected at baseline in the WHI population ^2^ and at exam 8 in the FOS.^3^

DNA methylation was assessed genome-wide with commercial arrays based on bisulfite conversion of unmethylated cytosines. In the REGICOR study, we used the Infinium MethylationEPIC BeadChip (Illumina, CA, USA). This array analyses over 850,000 CpGs per sample.^4^ After checking the DNA quality with Picogreen (Thermo Fisher Scientific, MA, USA), samples of REGICOR-1 were analysed in 13 batches of the Infinium MethylationEPIC BeadChip in the Genomics and Epigenomics Service of the Bellvitge Institute for Biomedical Research (Barcelona, Spain). Samples of REGICOR-2 were also analyzed at the same facilities, in three batches. In the WHI and FOS studies, the Infinium HumanMethylation450 BeadChip (Illumina, CA, USA) was used. This array analyses over 485,000 CpGs per sample,^5^ from which 439,562 are included in EPIC BeadChip.^4^ Assessment of DNA methylation for these two samples has been described elsewhere.^2,6^

**Quality control of DNA methylation data**

Quality control of the raw methylation data was previously described for the Infinium HumanMethylation450 BeadChip data by our group.^7^ This pipeline was applied to FOS and WHI.

*Quality control of the samples analysed with the Illumina MethylationEPIC BeadChip*

We removed the samples with a detection p-value >0.05 in at least 1% of the probes using the *pfilter* function of the *wateRmelon* R package available through the Bioconductor repository. We also discarded those samples that did not cluster in the corresponding sex cluster based on the DNA methylation levels in the X chromosome using the *methylumi* R package available through the Bioconductor repository.

*Quality control of the CpGs analysed with the Illumina MethylationEPIC BeadChip*

We excluded those probes with both a detection p-value >0.05 in at least 1% of the samples and a beadcount <3 in at least 5% of the samples using *wateRmelon* R package available through the Bioconductor repository. We further removed those probes reported by Illumina to be discarded due to underperformance (n=1,031) and changes in the manufacturing process (n=977). Finally, we excluded those probes corresponding to a methylation site different from a CpG site and those that could hybridize in more than one genomic region (n=43,979).^8,9^

**DNA methylation measurement**

Methylation status at each CpG site was reported by β-values, which are more intuitively interpreted than M-values.^10^ β-values range between 0 (completely unmethylated) and 1 (completely methylated).

They were calculated according to equation 1:

$\beta-value=\frac{M_{i}}{M_{i}+U_{i}+\alpha}$	Equation 1

Where:

-	M_i_ = intensity of methylated probe,

-	U_i_ = intensity of unmethylated probe, and

-	α = 100; constant offset.

To remove potential sources of technical variation not related to the underlying biology, we standardized the β-values by batch, as described previously by our group,^7^ using Equation 2:

$Z=\frac{(X-\bar{X})}{\sqrt{\frac{\sum{(X-\bar{X})}^{2}}{(n-1)}}}$	Equation 2

Where:

-	Z = standardized β-value,

-	X = β-value for a specific individual,

-	X̄ = mean of β-value for a specific batch, and

-	n = sample size.

Finally, to avoid the influence of extreme values, we excluded those CpGs with a β-value 4 standard deviations higher and lower from the mean.

**Genomic information**

We obtained genomic information of the CpGs using the manifest and annotation provided by Illumina and contained in the corresponding R packages available through the Bioconductor repository (*IlluminaHumanMethylation450kanno.ilmn12.hg19* and *IlluminaHumanMethylationEPICanno.ilm10b2.hg19*).

**Covariates assessment**

The REGICOR study specifically trained a group of nurses to collect blood samples and sociodemographic, lifestyle, and cardiovascular risk factors information using validated methods and questionnaires.^11^ Data from the FOS and the WHI sample were obtained through the Genotypes and Phenotypes database (http://dbgap.ncbi.nlm.nih.gov; project number #9047). The procedures used to collect data and blood samples of the REGICOR, WHI and the FOS populations were previously described.^2,12,13^ We considered the following covariates:

- *Smoking*: self-reported and categorized as current smokers (at least 1 cigarette/day or quitted smoking within the year before the visit) or non-smokers (never smoked or quit smoking at least one year before the visit),
- *Diabetes*: self-reported diabetes or treatment (REGICOR), glucose levels ≥126 mg/dL or treatment (WHI and FOS),
- *Hypercholesterolemia*: self-reported high cholesterol levels or treatment (REGICOR), total cholesterol ≥240 mg/dL or LDL-C ≥160 mg/dL or treatment (WHI and FOS),
- *Hypertension*: self-reported hypertension or treatment (REGICOR), SBP ≥140 mmHg or DBP ≥90 mmHg or treatment (WHI and FOS),
- *Estimated peripheral blood cell counts*,^14^
- *Two surrogate variables*.^15^

When applicable, we excluded from analysis those individuals with no information available regarding the cardiovascular risk factors.

**Statistical analysis**

*Analyses of the association between DNA methylation and cardiovascular outcomes*

A.- Main strategy: EWAS

The *bacon* R package controls for bias and inflation using a Bayesian method based on the estimation of the empirical null distribution and was used in previous EWAS.^16–18^ We used coefficients and standard errors from the regression models as the input data and set a random seed at 123.

B.- Complementary strategy: candidate gene analyses

We followed the same pipeline as in the epigenome-wide strategy, but in REGICOR-1 we performed the association analyses of the cardiovascular-related CpGs and the CpGs located in the genes differentially methylated in association with cardiovascular outcomes (Supplementary Table 1).^19–22^

*Assessment of the association between the identified CpGs and CVRFs*

Considering methylation as the outcome, we used linear regression models adjusted for age and sex in the case of the REGICOR and the Framingham populations, and for age and ethnicity in the WHI sample. In the case of the REGICOR samples, the continuous variables were only available for the control individuals. We meta-analysed the results from the four populations using a fixed-effects meta-analysis weighted by the inverse of the variance. The *p* value threshold was estimated as 0.05 divided by the multiplication of the number of CVRFs and the number of CpGs assessed.

*Methylation risk scores (MRS)*

The MRSs summarize an individual’s epigenetic predisposition to suffer a cardiovascular event. The weights for each CpG were based on the coefficients of the meta-analysis of Model 1 following Equation 3:

$MRS=\sum_{i=1}^{N} \beta_{i}^{meta}\cdot\beta_{i}^{meth}$	Equation 3

Where:

-	MRS = methylation risk score for a specific individual,

-	i = CpG,

-	N = CpG sample size,

-	β_meta_ = coefficient of the meta-analysis for each CpG, and

-	β_meth_ = standardized β-value for each CpG.

*Evaluation of the predictive capacity of CVR functions including the MRS*

The estimated risk for each individual was computed using Cox regression, according to Equation 4:

$CVR=1-S_{\bar{X}}^{\sum_{j=1}^{p} \beta_{j}^{F}\cdot(F_{j}-\bar{F}_{j})+\beta^{MRS}\cdot(MRS-\bar{MRS})}$	Equation 4

Where:

-	*1-S* = probability of presenting a CVD event in the next 10 years based on the incidence of CVD in the population,

-	*β^F^* = effect size of each CVRF,

-	*F_j_* = CVRFs per individual (logarithm of age, total cholesterol and HDL-C, and SBP (treated and not treated), sex, smoking status, hypertension treatment and diabetes),

-	$\bar{F}_{j}$= population mean of the CVRFs (logarithm of age, total cholesterol and HDL-C, and SBP (treated and not treated), sex, smoking status, hypertension treatment and diabetes),

-	*β^MRS^* = effect size of the MRS,

-	*MRS* = MRS per individual, and

-	$\bar{MRS}$= population mean of the CVRFs.

First, we evaluated the calibration of the models using the Hosmer-Lemeshow test.^23^ Next, we assessed the discriminative capacity of the models using the concordance index (c-statistic),^24^ applying the *rcorr.cens* function of the *Hmisc* R package. Last, we calculated the reclassification improvement using the net reclassification improvement (NRI) index^25^ with the *nricens* R package. In this regard, we defined three risk categories (low, intermediate and high) with cut-off points defined according to guidelines for 10-year risk reported by the NCEP Panel:^26^ [0–10)%, [10–20)%, ≥20%, respectively). We calculated the expected number of events at 5 years in each risk category (thus, [0–5)%, [5–10)%, ≥10%) using Kaplan-Meier estimates. Confidence intervals for the Kaplan-Meier estimates were obtained from the bootstrapping method applied by the *nricens* R function (*niter* = 1,000 bootstrap samples). We also analysed the NRI in the group of individuals with intermediate CVR (clinical NRI). To correct for bias in the NRI estimation among individuals with intermediate risk, we used the method proposed by Paynter and Cook.^27^

**Analysis of the causality of associations between DNA methylation and cardiovascular outcomes**

To run the mendelian randomization analyses, we selected the following methods as options: (a) clumping to prune SNPs for linkage disequilibrium (LD); (b) proxy SNPs through LD tagging (minimum LD R^2^=0.8) if one SNP is not present in an outcome dataset, allowing palindromic SNPs (MAF threshold for alignment=0.3); (c) alignment of strands for palindromic SNPs for allele harmonization (i.e. effects of the SNPs on DNA methylation and on the outcome correspond to the same allele); and (d) Wald ratio, maximum likelihood, MR Egger, weighted median, Inverse variance weighted, Inverse variance weighted (fixed effects), and weighted mode.

**SUPPLEMENTARY FIGURES**

**Supplementary Figure 1.** Manhattan plots of the associations between DNA methylation and acute myocardial infarction in the discovery stage (REGICOR-1). Model 1 was adjusted for estimated cell counts and two surrogate variables. Model 2 was further adjusted for smoking status. Model 3 was additionally adjusted for diabetes, hyperlipidaemia and hypertension. Inflation was corrected using the bacon R package; plots are given before (left) and after (right) the correction.

| **Model 1** | **Non-corrected** | **Corrected** |
| --- | --- | --- |
|  | 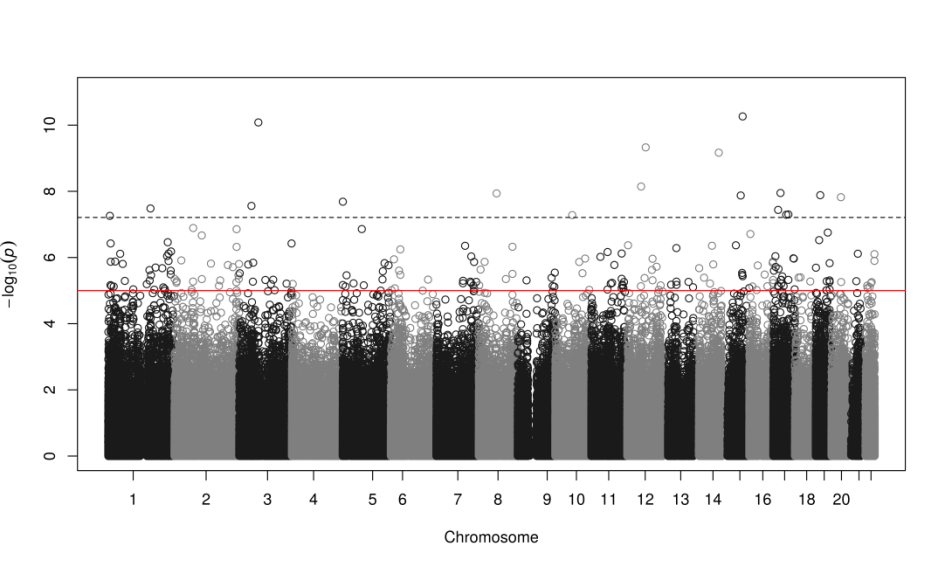 | 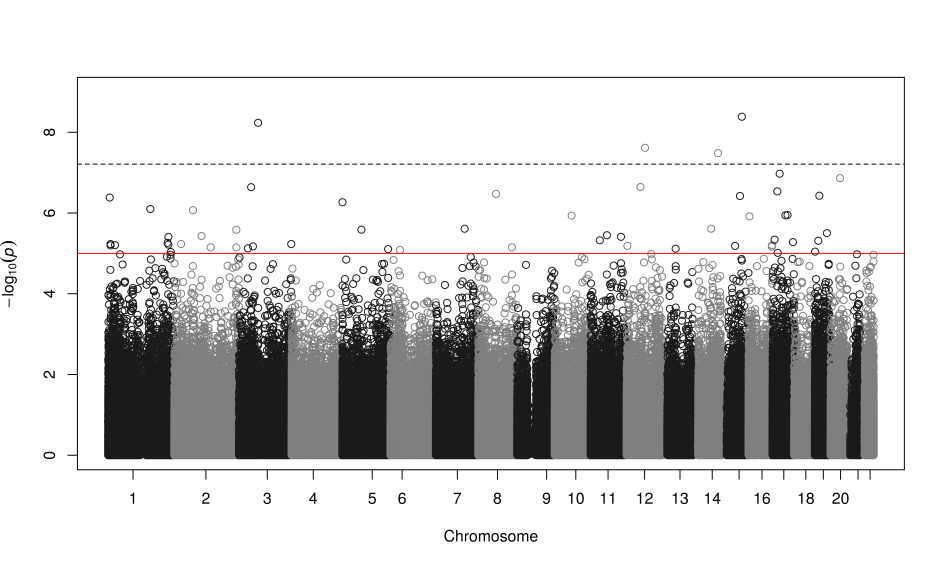 |

|  | **Non-corrected** | **Corrected** |
| --- | --- | --- |
| **Model 2** | 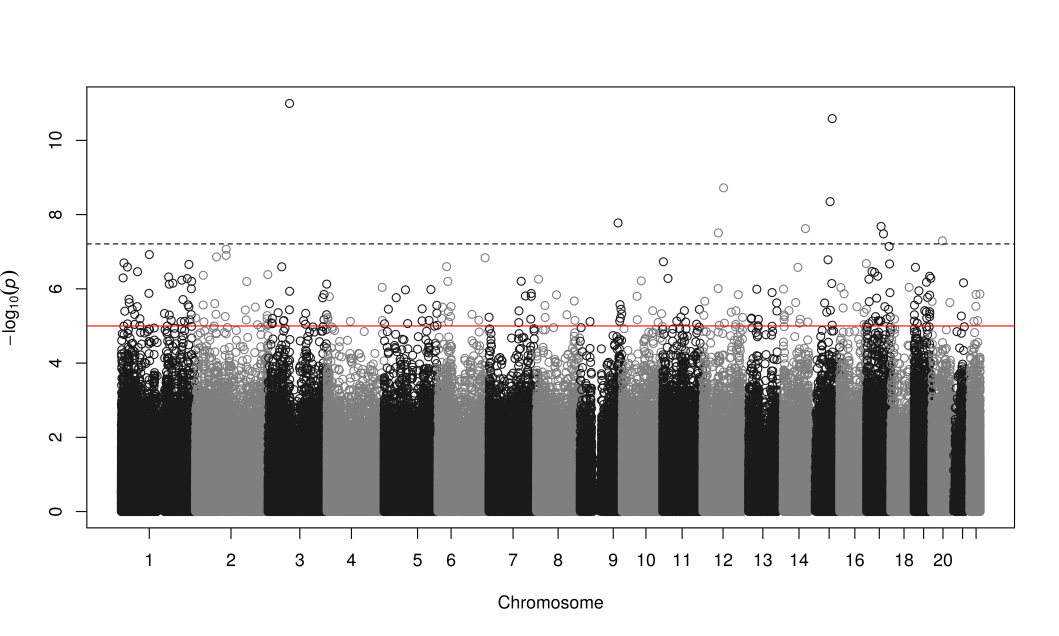 | 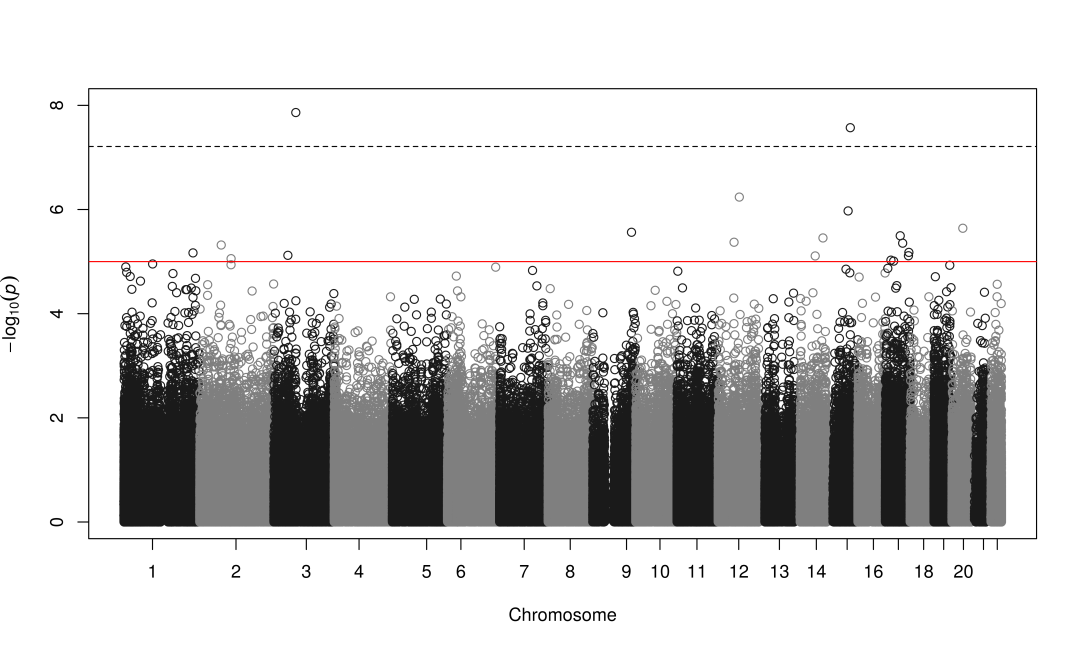 |
|  | **Non-corrected** | **Corrected** |
| **Model 3** | 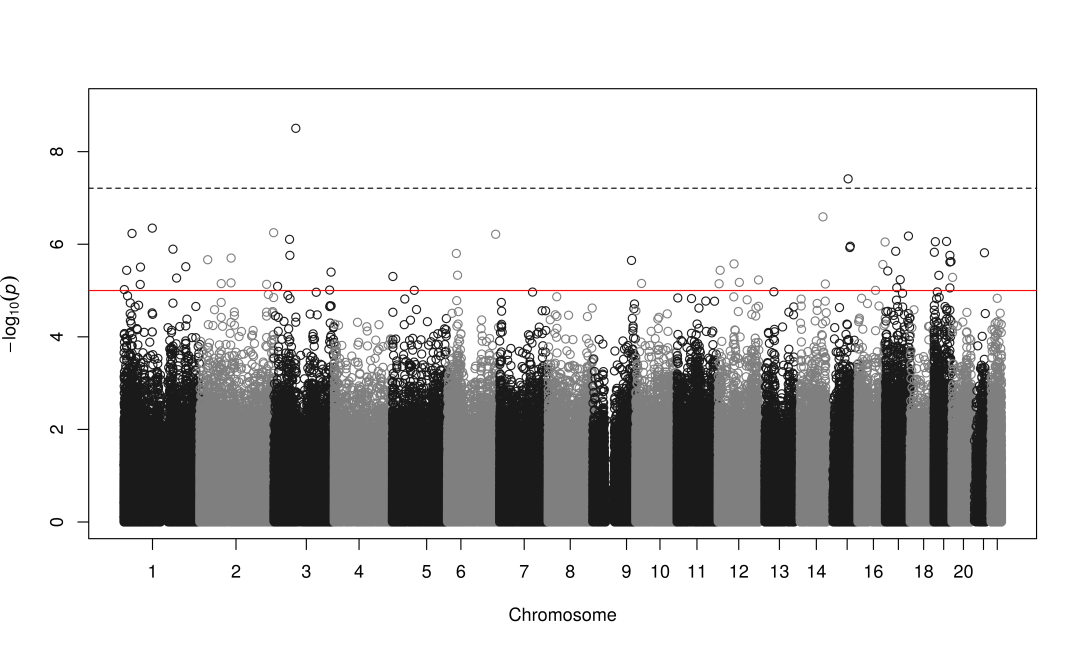 | 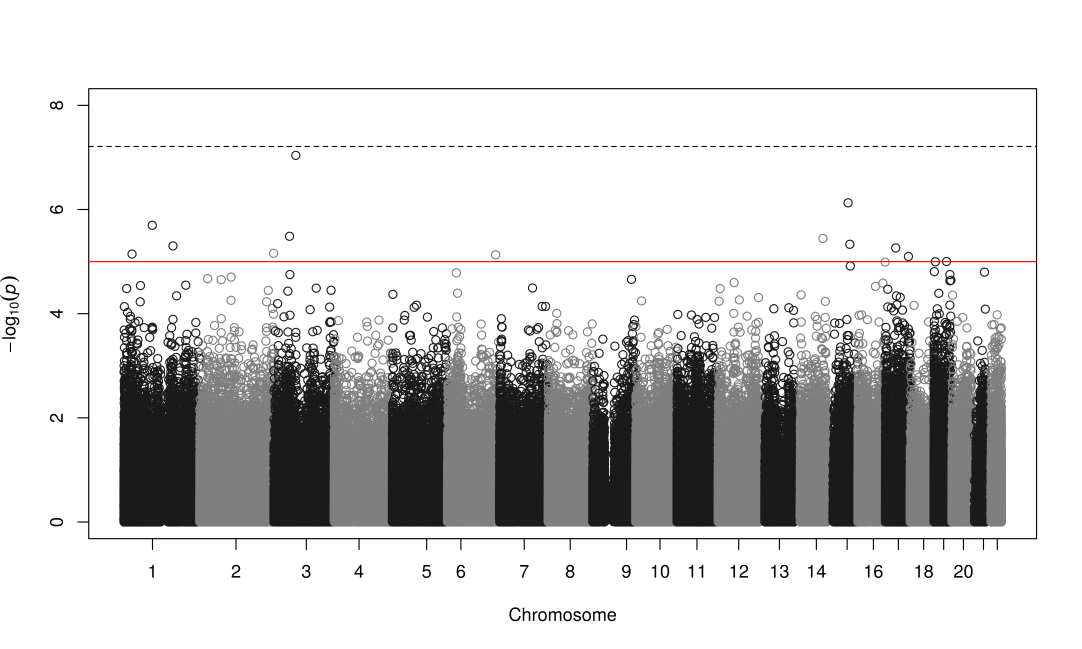 |

**Supplementary Figure 2.** QQ plots of the associations between DNA methylation and acute myocardial infarction in the discovery stage (REGICOR-1). Model 1 was adjusted for estimated cell counts and two surrogate variables. Model 2 was further adjusted for smoking status. Model 3 was additionally adjusted for diabetes, hyperlipidaemia and hypertension. Inflation was corrected using the bacon R package; plots are given before (upper row) and after (bottom row) the correction.

|  | **Model 1** | **Model 2** | **Model 3** |
| --- | --- | --- | --- |
| **Non-corrected** | 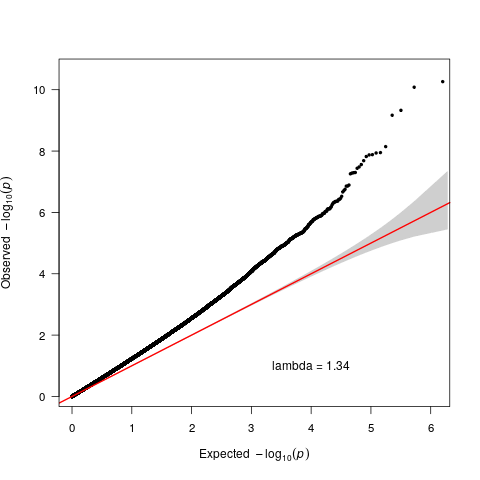 | 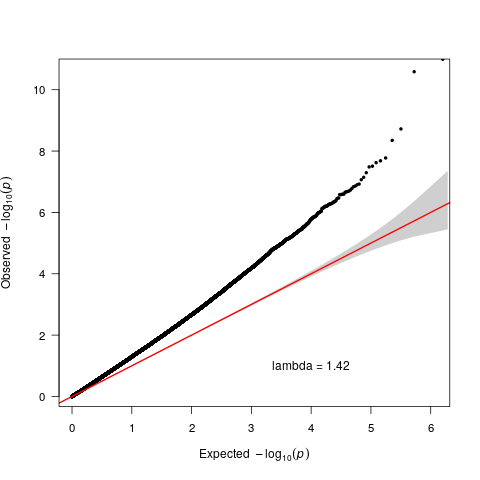 | 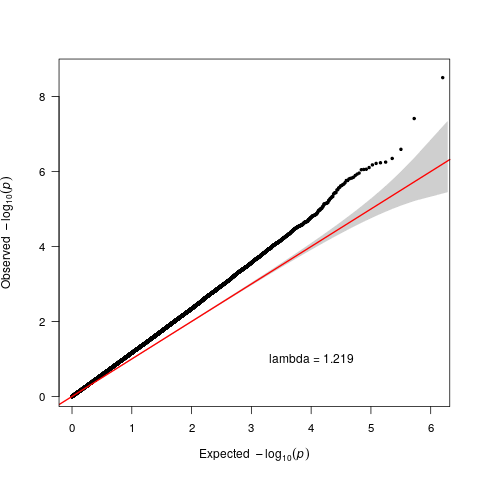 |
| **Corrected** | 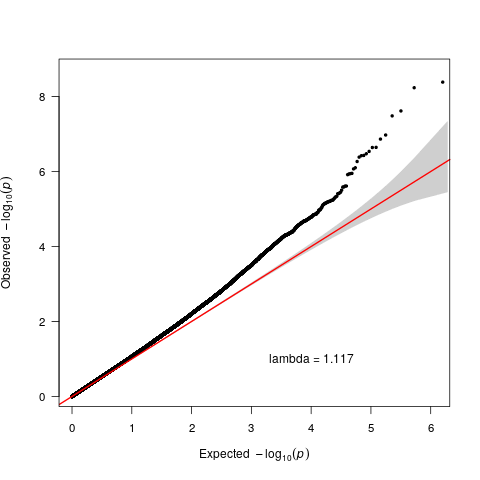 | 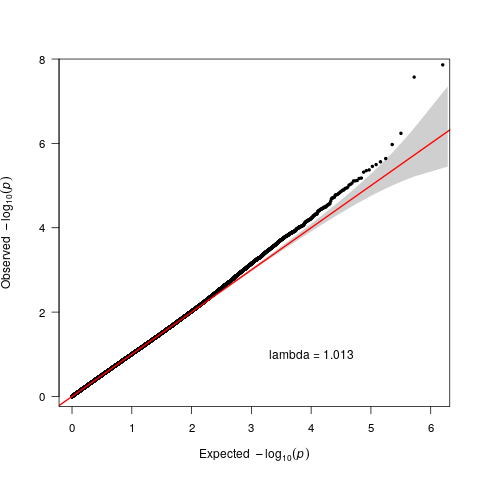 | 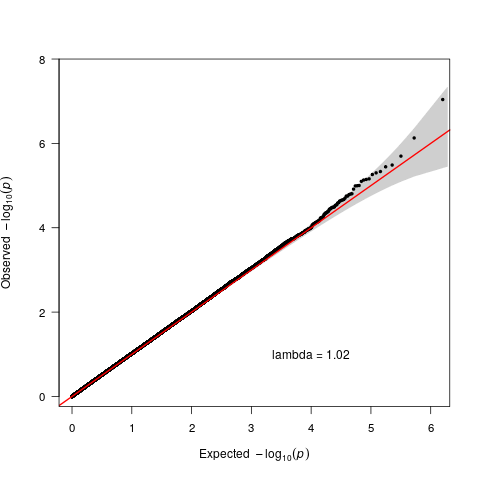 |

**Supplementary Figure 3. Venn diagram showing the overlap of CpGs found in the three models.** The number of CpGs instead of their identifiers is given.

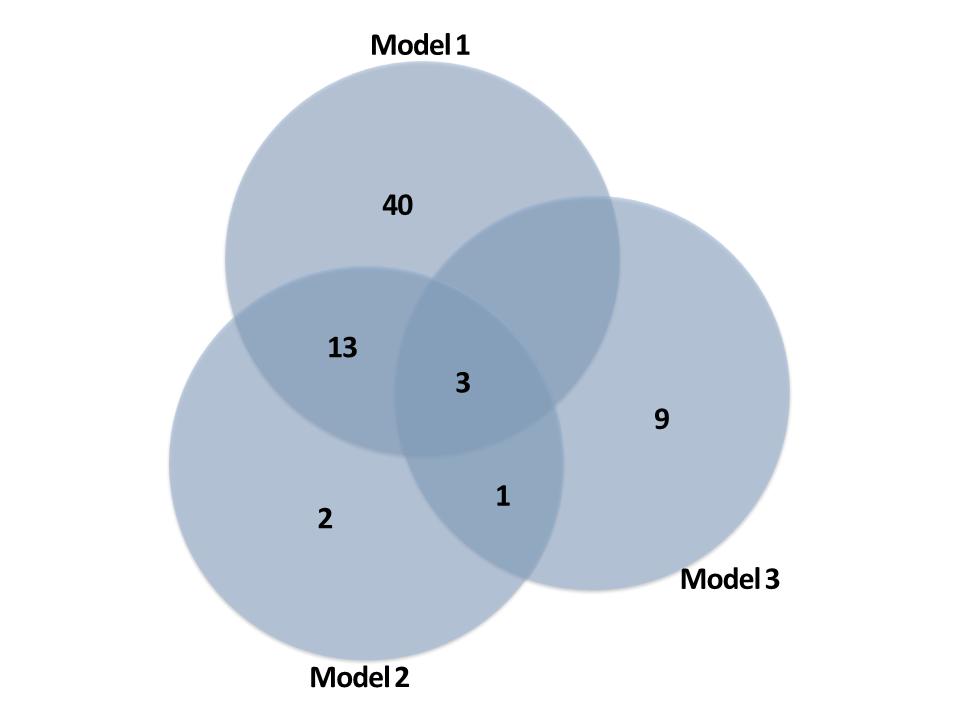

**Supplementary Figure 4. Ingenuity pathway analysis: Functional classification of the genes identified as differentially methylated in association with myocardial infarction.** “Diseases and functions” significant after the Benjamini-Hochberg multiple testing correction are included. A) Molecular and cellular functions; B) Physiological system development and functions; and C) Diseases and disorders.

*A) Molecular and cellular functions*

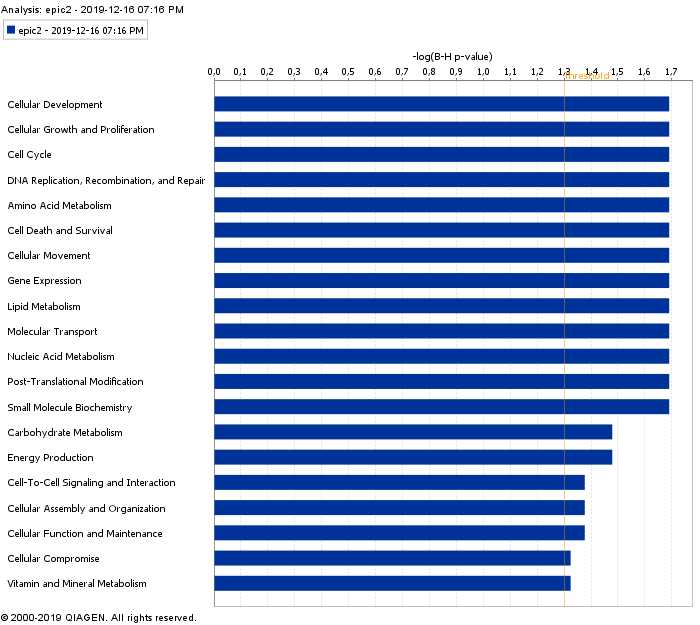

*B) Physiological system development and functions*

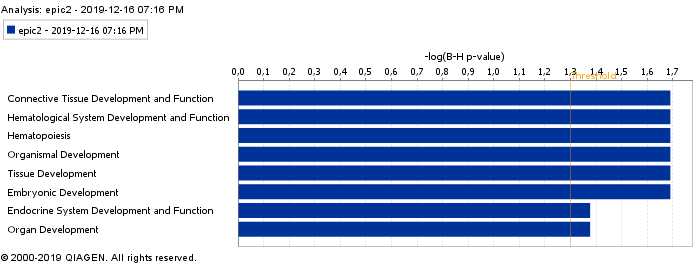

*C) Diseases and disorders*

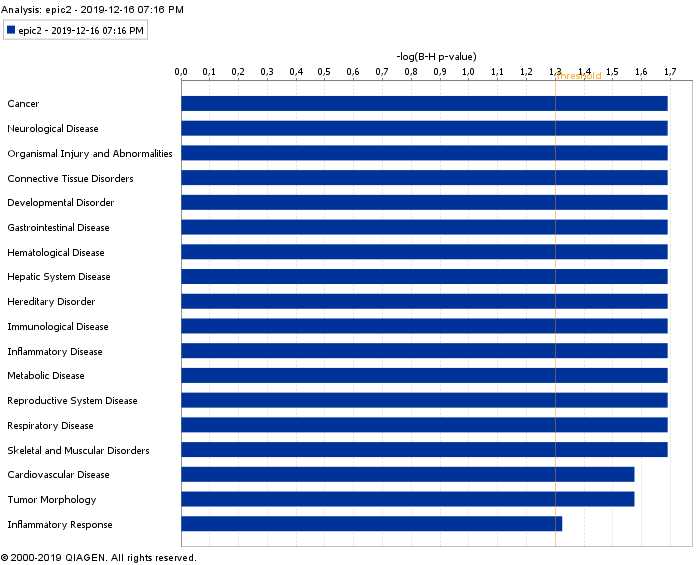
